# Arena3D^web^: Interactive 3D visualization of multilayered networks

**DOI:** 10.1101/2020.11.20.391318

**Authors:** Evangelos Karatzas, Fotis A. Baltoumas, Nikolaos A. Panayiotou, Reinhard Schneider, Georgios A. Pavlopoulos

**Author notes:** Present Address: Georgios A. Pavlopoulos, Institute for Fundamental Biomedical Research, BSRC “Alexander Fleming”, 34 Fleming Street, Vari, 16672, Greece.

## Abstract

Efficient integration and visualization of heterogeneous biomedical information in a single view is a key challenge. In this study, we present Arena3D^web^, the first, fully interactive and dependency-free, web application which allows the visualization of multilayered graphs in 3D space. With Arena3D^web^, users can integrate multiple networks in a single view along with their intra- and inter-layer connections. For clearer and more informative views, users can choose between a plethora of layout algorithms and apply them on a set of selected layers either individually or in combination. Users can align networks and highlight node topological features, whereas each layer as well as the whole scene can be translated, rotated and scaled in 3D space. User-selected edge colors can be used to highlight important paths, while node positioning, coloring and resizing can be adjusted on-the-fly. In its current version, Arena3D^web^ supports weighted and unweighted undirected graphs and is written in R, Shiny and JavaScript. We demonstrate the functionality of Arena3D^web^ using two different use-case scenarios; one regarding drug repurposing for SARS-CoV-2 and one related to GPCR signaling pathways implicated in melanoma. Arena3D^web^ is available at http://bib.fleming.gr:3838/Arena3D.

## INTRODUCTION

Simple networks are often used to model relationships between entities in real-world systems. Typical examples of such networks in the biomedical field are the gene co-expression networks, sequence similarity networks (SSNs), protein-protein interaction networks (PPIs), metabolic networks, signal transduction networks, gene regulatory networks, gene-disease networks, evolution networks, bipartite graphs, food webs and others (1). Most of these networks come with certain topological features and fall into one of the Erdos–Rényi (random), Watts-Strogatz (small-word) and Barabási–Albert (scale-free) categories (2). Efficient visualizers take advantage of these topological properties and utilize advanced graph-drawing (layout) algorithms (e.g. force-directed, hierarchical, orthogonal and spectral) to generate informative and appealing 2D/3D views thus minimizing any possible node overlaps and connection crossovers (3). Among a decent variety of applications today (4, 5), widely-used interactive 2D visualizers include the Cytoscape (6), Cytoscape.js (7), String (8), NORMA (9), Gephi (10), Pajek (11) and Tulip (12) whereas successful 3D visualizers include the Graphia (Kajeka) and BioLayout Express (13).

Despite the great functionality and interactivity of these viewers, most of them currently offer over-simplified views and often fail to capture patterns in today’s data heterogeneity and complexity. In the biomedical field for example, a new trend is to integrate heterogeneous information from various repositories in a single view or support experimental findings (e.g. biomarker detection) in multiple ways (e.g. genomics, proteomics, metabolomics). Therefore, visualizers able to offer a more structured analysis along with contemporary multilayer network visualization are necessary (14). While this area of network analysis is still in its infancy, few implementations in this direction have been already proposed. The Arena3D standalone version (15, 16) for example, enables an interactive multilayered graph visualization but is poor in the inter/intra-layer layouts that it offers. Pymnet is a python library which can produce high quality static images of multilayer and multiplex networks and therefore, familiarity with Python programming language is required. Similarly, Py3plex (17) a python library, which handles multilayered networks and enables common operations such as aggregation, slicing, indexing and traversal but lacks interactivity and fails to solve the edge overlapping problem. Mully (18) is an R package to create, modify and visualize multilayered graphs but lacks interactivity too. Finally, MuxViz (19) is a framework which is more suitable for layer analysis and visualization of geospatial data.

In this article, we propose Arena3D^web^, the first web application which offers a fully interactive 3D visualization and analysis of multilayered networks. Arena3D^web^ comes with many inter/intra-layer layouts and network metrics for node scaling, is dependency-free and is easily accessible to non-experts from a web browser. It is written in R, Shiny and JavaScript whereas the backend calculations are based on the igraph library (20). We believe that Arena3D^web^ is a general-purpose tool which can successfully address many of the complexity issues in the biomedical field and can offer informative and appealing visualizations suitable for knowledge integration, representation, transfer and communication.

## MATERIAL AND METHODS

### Input and export files

In its current version, Arena3D^web^ mainly accepts three different types of tab-delimited files as input. The main file is a four- or a five-column file describing the network in a simple format like: *node i from layer x is connected to node j from layer y*. In the case of a weighted graph, weights can be added as an extra column. In addition to the main file, users are allowed to upload two more files with further information about the node and edge attributes. In the first case, nodes can be accompanied by a URL, a description, a color, and a custom size whereas in the second case, edges can be accompanied by a color. For convenience, the column order in any of the aforementioned files is insignificant as long as headers are used strictly. Notably, users can save and export the status of a network at any time in an Arena3D^web^ file format. This file contains all the necessary information from nodes and connections to colors and 3D coordinates. It is a text file format which users can modify manually at any time.

### Graph drawing and layouts

As a default option, Arena3D^web^ places all uploaded layers in parallel and spreads them horizontally. Once the main network has been loaded and rendered, users can choose between a variety of layout algorithms to adjust the node coordinates. By applying a layout algorithm on a selected layer or a set of selected layers individually, users can eliminate the intra-layer line crossovers without impacting the inter-layer connections. On the contrary, when users decide to apply a layout algorithm on a set of selected layers in combination, then more focus is given on eliminating the inter-layer crossovers. In the second case, the layout algorithm will handle all nodes from the selected layers and their connections as one unified network and will place them back on their originating layers once the layout algorithm has converged. The latter is a powerful feature for creating network pseudo-alignments (Figure 1, layers 5 and and generating appealing and informative views. Finally, users are allowed to apply any of the offered layout algorithms locally on a set of selected nodes per layer.

**Figure 1.**
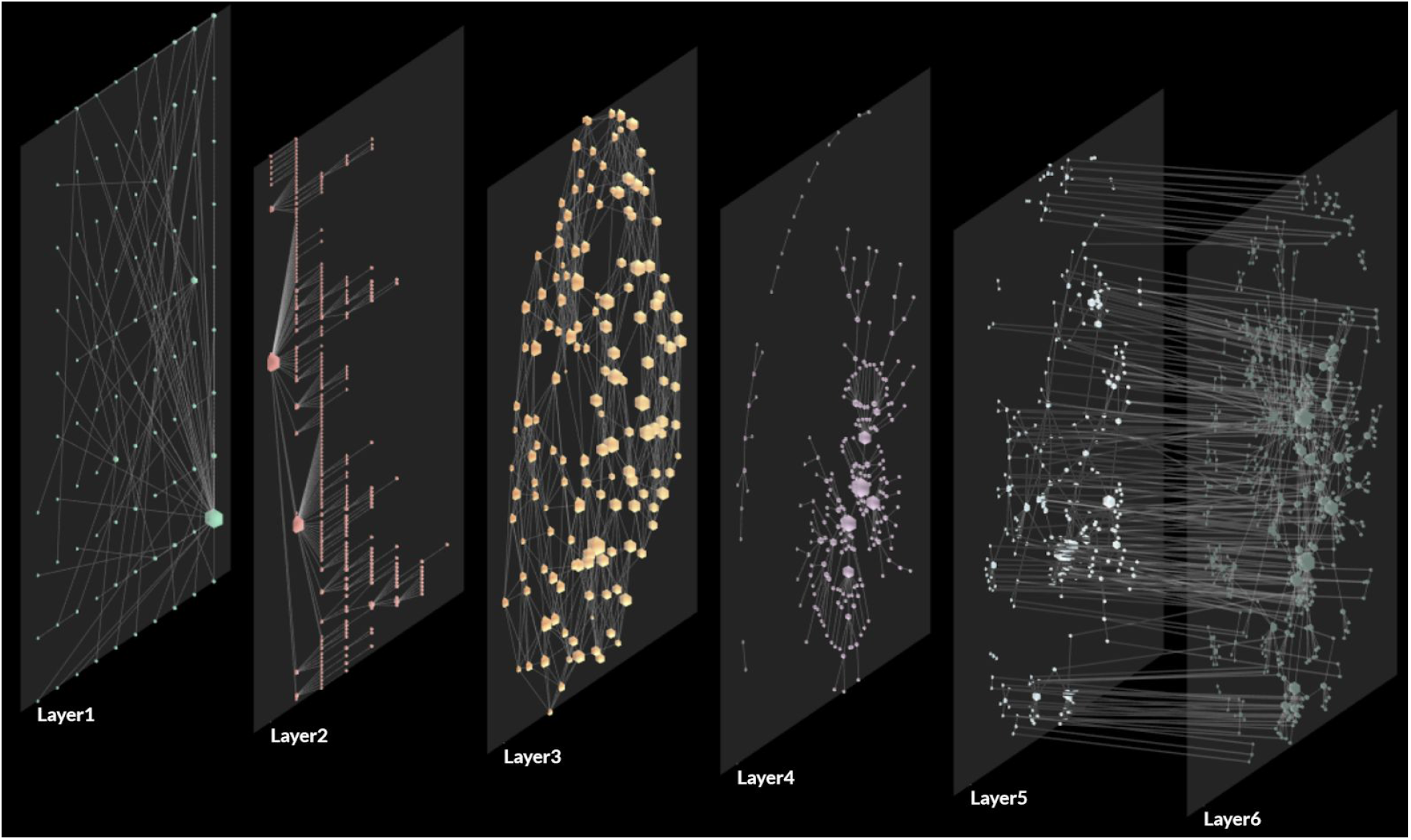
Random networks with different topologies were drawn with the use of various layout algorithms supported by igraph and subsequently by Arena3D^web^. Nodes on layer 1 are placed on a grid. Nodes on layer 2 have been placed in a hierarchy using the Reingold-Tilford layout algorithm. Nodes on the first two layers are scaled based on degree. Layer 3 shows a small-world network drawn using the Kamada-Kawai layout algorithm. Layer 4 shows a scale-free network also drawn with the Kamada-Kawai layout algorithm. Nodes on layers 3 and 4 are scaled based on their clustering coefficient. Layers 5 and 6 have been combined and are drawn using the Fruchterman-Reingold layout algorithm. Both layers were handled as a unified network and nodes were placed back to their layers after the completion of the layout algorithm, thus giving a sense of a network alignment. The node scale of these two layers is relative to their degree.

In this version, Arena3D^web^ supports a variety of layout algorithms implemented as part of the R/igraph library (20). Briefly, these are:

- *Circle*: It is a simple layout and places the vertices on a circle, ordered by their vertex ids.
- *Grid*: This simple layout places vertices on a rectangular 2D grid.
- *Random*: This function places the vertices of the graph on a 2D plane uniformly using random coordinates.
- *Star*: It places the vertices of a graph on the plane, according to the simulated annealing algorithm developed by Davidson and Harel. It is a simple layout generator that places one vertex in the center of a circle and the rest of the vertices equidistantly on the perimeter.
- *Reingold-Tilford* (21): It is a tree-like layout more suitable for trees, hierarchies and graphs without many cycles.
- *Sugiyama*: Like Reingold-Tilford, this layout algorithm is more suitable for hierarchies and layered directed acyclic graphs.
- *Fruchterman-Reingold* (22): It places nodes on the plane using the force-directed layout algorithm developed by Fruchterman and Reingold.
- *Davidson-Harel* (23): It is a force-directed algorithm which uses simulated annealing and a sophisticated energy function to place nodes on a plane.
- Distributed Recursive (Graph) Layout (24): DrL is a force-directed graph layout toolbox focused on real-world large-scale graphs.
- *Multidimensional scaling* (25): It aims to place points from a higher dimensional space on a 2D plane, so that the distance between the points are kept as much as possible.
- *Kamada-Kawai* (26): This force-directed layout places the vertices on a 2D plane by simulating a physical model of springs.
- *Large Graph Layout (LGL)*: A force-directed layout suitable for larger graphs.
- *Graphopt*: A force-directed layout algorithm, which scales relatively well to large graphs.
- *Gem* (27): It places vertices on the plane using the GEM force-directed layout algorithm.

In Figure 1, we show representative views which demonstrate some of the aforementioned functionalities. Nodes on the first layer have been placed on a simple grid whereas node coordinates on the second layer (random acyclic graph) have been generated by the Reingold-Tilford layout. The force-directed Kamada-Kawai layout algorithm has been applied separately on a small-world and a scale-free random network on layers 3 and 4 accordingly. Finally networks on layer 5 and 6 have been treated as one unified network aligned with the use of the Fruchterman-Reingold force-directed algorithm.

### Topological features

Nodes on a set of selected layers can be resized according to topological properties such as the *degree*, *clustering coefficient* and *betweenness centrality*. For example, nodes with higher connectivity degrees can be forced to appear larger. This function can be performed on each of the selected layers individually or in combination. In the first case, only the intra-layer connections are taken into account whereas in the second case both the intra- and inter-layer connections are considered. Briefly, as *degree deg*_*i*_, we define the total number of connections adjacent to a node *i*. The *clustering coefficient* of a node *i* shows whether this node has the tendency to form clusters and is defined as the number of edges between a node’s neighbors divided by the number of all possible connections between these neighbors. The *betweenness centrality* highlights nodes which can act as mediators in order for two communities to communicate with each other. It is calculated as 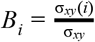 where σ_*xy*_ is the total number of shortest paths from node *x* to node *y* and σ_*xy*_ (*i*) is the number of those paths that pass through the node *i*. For demonstration purposes, in Figure 1, nodes on layers 1 and 2 have been resized based on their connectivity degree. Nodes on layers 3 and 4 have been resized based on their clustering coefficient. Finally, nodes on layers 5 and 6 have been resized based on their degree, while considering both layers as one unified network.

### Navigation controls

Arena3D^web^ comes with various control buttons for easy 3D navigation (Figure 2). These buttons are grouped in three main categories. These are: *i)* buttons for the whole scene, *ii)* buttons for each layer, *iii)* buttons for the nodes within a layer.

**Figure 2.**
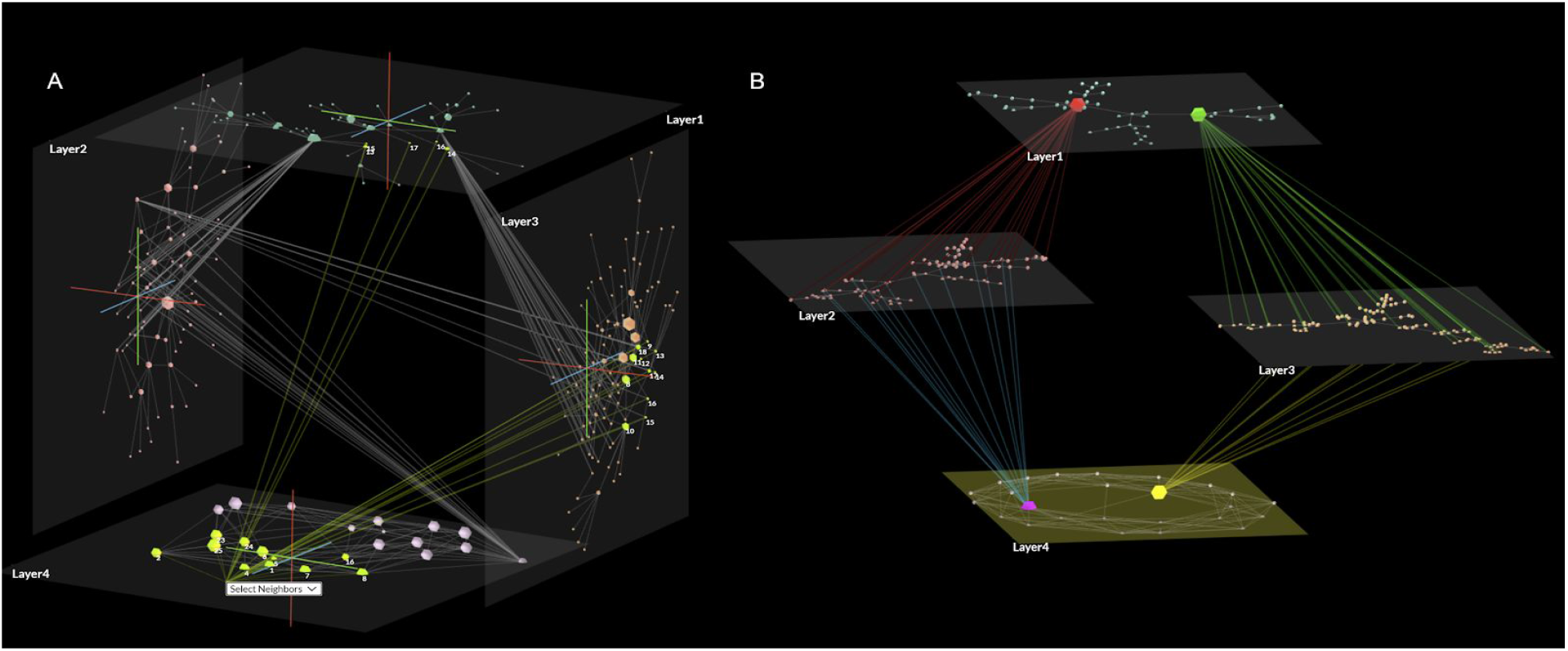
Random networks visualized by Arena3D^web^. A) Layers 1-4 are placed in a 3D cube. Node sizes on layers 1 and 2 are adjusted by connectivity degree, nodes on layer 3 by betweenness centrality and nodes on layer 4 by clustering coefficient. The Kamada Kawai layout algorithm has been applied on each layer individually. First neighbors of a node and their connections are colored yellow. B) Layers 1-3 host three different scale-free random networks whereas layer 4 hosts a small-world random network. Based on a node attributes file, nodes with higher connectivity degree are resized and colored red, green, purple and yellow. Based on an edge attributes file, the inter-layer edges of the upscaled nodes are colored red, green, blue and yellow. The Fruchterman-Reingold layout algorithm has been applied on each layer individually.

#### The Scene

Arena3D^web^ uses a global cartesian coordinate system and adopts the red green and blue colors to highlight the x, y and z axes accordingly. The whole network can be rotated clockwise and counter-clockwise around any on the three axes using the navigation buttons. The rotation angle and subsequently the rotation speed can be adjusted manually. Zooming, panning and orbiting are controlled by mouse events as well as by keyboard strokes.

#### The Layer

In Arena3D^web^, each layer comes with its own individual coordinate system. Users are allowed to rotate and translate any layer around any of the x, y, z axes and place them anywhere in 3D space. In the case where multiple layers have been selected, each transformation will be applied to each of them individually. Like before, the rotation angle as well as the translation step can be adjusted manually. In addition, each layer floor can be scaled up or down by a user-defined factor and node coordinates on the corresponding layer will be adjusted automatically. Finally, to avoid incomprehensive views when layers are placed too close to each other, users have the option to expand or collapse the view by adjusting their in-between distance.

#### The nodes

Nodes can be placed anywhere on a plane regardless the position and orientation of the layer. Users can select a single node or multiple nodes simultaneously using a rectangle selection and translate them in 3D space. To cope with node and label overlapping problems, users can apply any of the offered layout algorithms locally on a set of selected nodes or increase and decrease their local coordinates thus giving the feeling of repulsion or attraction. Like before, the translation step can be manually adjusted. Finally, node sizes can be scaled manually with the use of a slide bar.

### Styles and colors

Arena3D^web^ comes with a plethora of options to avoid incomprehensive visualizations. Users can show and hide the labels of each layer as well as the layer itself along with the corresponding connections. Similarly, users can show and hide the coordinate system of the whole scene as well as the coordinate systems of each individual layer. Nodes and edges can be manually colored to highlight important paths whereas edge weights are represented by an edge’s opacity. The lower the weight, the more transparent the edge’s color and vice versa. The background color as well as each layer’s floor color can be changed with the use of a color picker whereas each floor’s transparency can be adjusted manually. Finally, the inter-layer and the intra-layer connection opacities can be adjusted. This way, one can emphasize on the intra- or the inter-layer edges separately. Finally, Arena3D^web^ gives the option to highlight the first-neighbors of a node along with their corresponding connections or highlight the paths that this node is involved in across all layers.

### Implementation

Arena3D^web^ is mainly written in R and JavaScript programming languages. It’s frontend is written using the R/Shiny package, HTML, CSS and JavaScript. The Shiny package was used as the mediator to establish the communication between the R and JavaScript variables and functions. In the backend, the igraph library was used for all network layout algorithms as well as for the calculation of the topological metrics used for node scaling. 3D graphics in the main view were handled by the three.js JavaScript library. The library uses the WebGL JavaScript API for graphics and 3D object rendering.

## RESULTS

In order to demonstrate how Arena3D^web^ can visualize heterogeneous complex information in real case scenarios, we provide two examples, the one related to drug repurposing for SARS-CoV-2 (network with 456 nodes and 985 edges) and the other related to GPCR signaling pathways implicated in melanoma (network with 64 nodes and 332 edges).

### Drug repurposing for SARS-CoV-2 - A case study

SARS-CoV-2 is the causative coronavirus for the disease COVID-19 (28) which has infected over 46 million people and has led to more than 1.2 million deaths (worldometers.info/coronavirus - 11/2020). In a recent study of Gordon et. al (29), 26 SARS-CoV-2 proteins have been cloned, tagged and expressed in HEK-293T/17 human cells in order to identify physically interacting proteins with the use of affinity-purification mass spectrometry (AP-MS) technology. Experiments led to the identification of 332 high-confidence protein–protein interactions between the SARS-CoV-2 virus and the human proteins. In addition to this, the study highlights existing drugs which target the human proteins and which can be potentially repurposed and be used for the COVID-19 treatment.

In more detail, the 2019-nCoV/USA-WA1/2020 SARS-CoV-2 genome (MN985325) was taken from the GenBank database (30) and was analyzed across its 29 possible open-reading frames (ORFs). The analysis reported 26 annotated protein baits which were found to interact with proteins in human cells. In many cases, these human proteins were found to form protein complexes, reported in the CORUM database (31). In addition, functional enrichment analysis reported Gene Ontology (GO) biological process terms (32), collected from the Molecular Signature Database (33). Interacting drugs with both viral and human proteins were identified through literature searching for targets and pathways, as well as based on cheminformatics searches in the 2020 IUPHAR/BPS Guide to Pharmacology (34) and the ChEMBL25 (35) databases.

In order to combine the heterogeneous information from this example in a single view, based on the data provided by the Gordon et al. study, we constructed a five-level multilayered network with Arena3D^web^ (Figure 3). In detail, these layers are: (i) human proteins and their interactions, (ii) protein complexes these proteins belong to, (iii) biological processes these proteins are involved in, (iv) SARS-CoV-2 proteins and their interactions with the human proteins and (v) possibly repurposable drugs and their respective targets in both the SARS-CoV-2 and human protein layers. To create a more comprehensive view, we treated the different layers as one network and aligned them via the Kamada-Kawai layout. Drugs on the last layer were grouped using a local circular layout and FDA-approved drugs were marked in a cerulean color.

**Figure 3.**
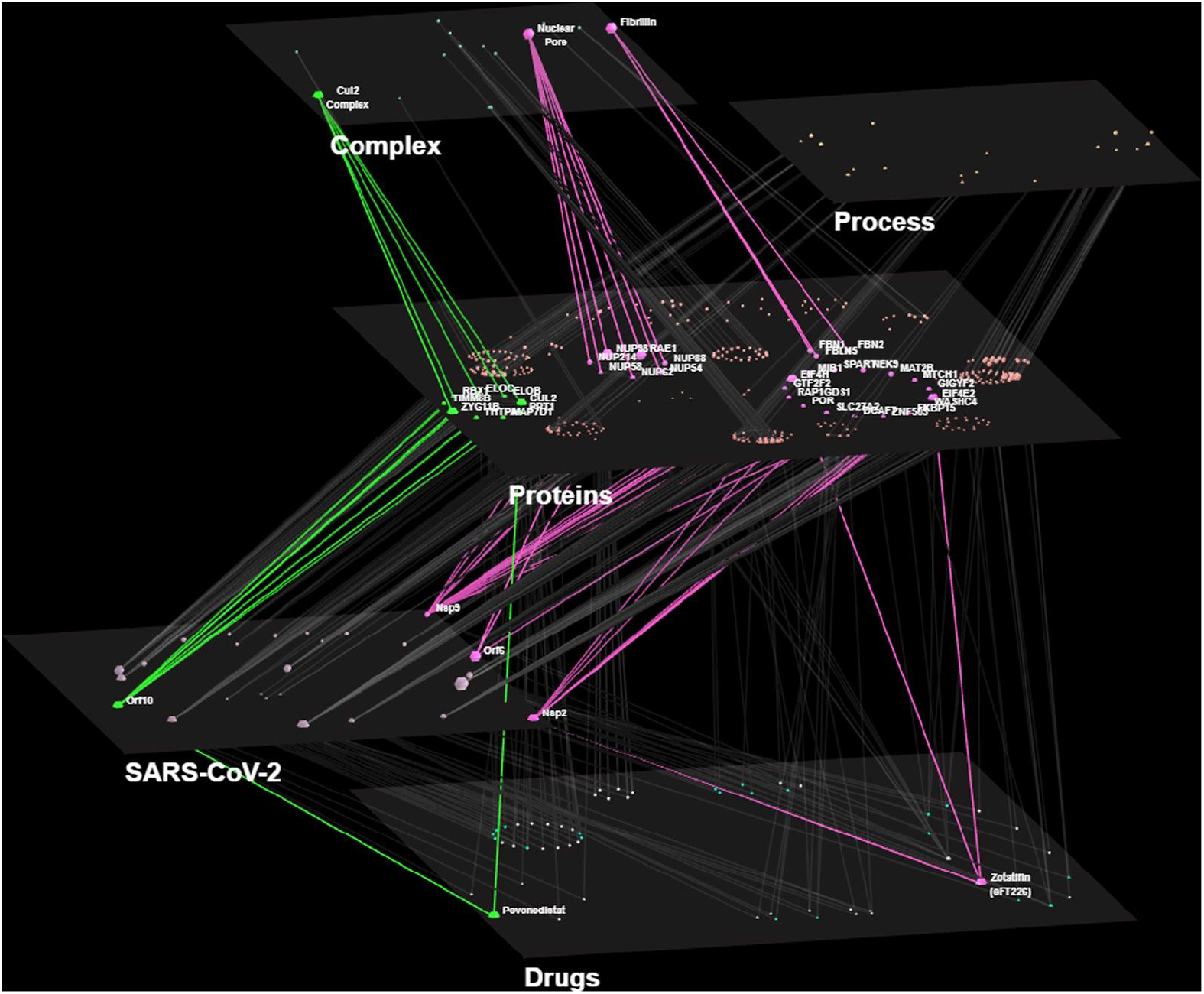
A multilayered COVID-19 related example based on the work of Gordon et al. Nodes are placed on five different layers (SARS-CoV-2 proteins, drugs, human proteins, protein complexes, biological processes). The vertices are aligned across every layer via the Kamada-Kawai layout algorithm. Two of the most interesting multilayer paths of this study are highlighted in green and pink colors respectively. With green color, pevonedistat directly targets the CUL2 protein complex and the SARS-CoV-2 ORF10 protein. With pink color, zotatifin directly targets eIF4 human proteins and the SARS-CoV-2 NSP2 protein. The latter indirectly interacts both with protein members of the nuclear pore and fibrillin complexes as well as with SARS-CoV-2 NSP9 and ORF6 proteins.

We focused on pevonedistat and zotatifin drugs as they target several SARV-CoV-2 proteins as well as certain human protein complexes. We highlight the inter-layer paths these drugs belong to with green and pink colors respectively as well as their protein targets and their distinct complexes. For better visualization, on the human protein layer, we applied the local circular layout where necessary.

With an at-a-glance view, we observe that the SARS-CoV-2 ORF10 ubiquitin ligase interacts with members from the cullin-2 (CUL2) RING E3 ligase complex and more specifically with the CUL2^ZYG11B^ complex. Either ORF10 binds to the CUL2^ZYG11B^ complex and hijacks it for ubiquitination and degradation of restriction factors, or ZYG11B binds to the N-terminal glycine in ORF10 and targets it for degradation (36). Pevonedistat is a NEDD8-activating enzyme (37) that allows the neddylation of CUL2, a process which is required for transfering ubiquitin to substrates (38). Therefore, pevonedistat may inhibit the ubiquitination of host proteins that is caused when ORF10 hijacks the CUL2^ZYG11B^ complex.

In general, coronaviruses produce their proteins via cap-dependent mRNA translation (39). Eukaryotic translation initiation factors 4 (eIF4) are proteins that, as their name implies, act as initiation factors for the translational process (40). Key eIF4F–cap binding components are attractive candidates for potential coronavirus treatments (41, 42). Zotatifin directly targets the SARS-CoV-2 NSP2 protein as well as the eIF4H and eIF4E2 human proteins. Specifically, Gordon et al. observed a strong antiviral effect through zotatifin, which inhibits eIF4A, a partner of eIF4H. Through the eIF4A inhibition, zotatifin may prevent the translation of a viral 5’ region by impeding its unwinding. The eIF4H human protein also interacts with the viral protein NSP9. Subsequently, NSP9 interacts with human proteins that belong to the nuclear pore and fibrillin protein complexes. Furthermore, the nuclear pore NUP98–RAE1 sub-complex was found to interact with the SARS-CoV-2 ORF6 protein. ORF6 perturbs nuclear transport in order to antagonize host interferon signalling in SARS-CoV (43). In SARS-CoV-2, the NUP98–RAE1-ORF6 interaction may have a similar effect.

Pevonedistat and zotatifin both target key human proteins that interact with SARS-CoV-2 proteins, as well as viral proteins which in turn interact with distinct human protein complexes. Since there are no overlaps among the targets of the aforementioned drugs, a cotherapy of the two should be further evaluated as a potential treatment against symptoms of COVID-19.

### GPCR signaling pathways implicated in melanoma - A case study

G-protein coupled receptors (GPCRs) are one of the largest and most diverse superfamilies of cell-surface receptors in eukaryotic organisms. They regulate the majority of cell responses related to signaling stimuli and have been implicated in a wide range of diseases. As a result, today GPCRs are targets for more than 30% of drugs on the market (44). Most GPCR functions are conducted through heterotrimeric G-proteins, composed by Gα subunits and Gβγ heterodimers, which act as molecular switches that modulate intracellular signaling cascades as responses to receptor activation (45). G-proteins are grouped, based on the Gα subunit, in four families, Gα_s_, Gα_i/o_, Gα_q/11_ and Gα_12/13_. They regulate the function of a wide and diverse range of effector proteins, including several enzymes, regulators of protein function, chaperones, transcription factors and cytoskeletal components. However, other signaling pathways also exist, either complementary to or independent from G-protein signaling. Phosphorylation of GPCRs by a special class of kinases, known as G-protein coupled receptor kinases (GRKs), leads to the internalization of the receptors; the internalized GPCRs can subsequently initiate signaling pathways regulating the expression of genes that control cell growth and/or apoptosis (46). Another example is GPCR oligomerization; a phenomenon during which multiple GPCRs form supramolecular homo- or hetero-complexes, leading to diverse signal transduction mechanisms (47). Finally, several GPCRs have been found to directly interact with effectors such as ion channels. All of the aforementioned signaling mechanisms result in complex and diverse signaling networks for GPCRs, with a single receptor capable of regulating a large number of pathways and therefore, a wide range of cell functions (48).

GPCR signaling can be deregulated in cancer; either due to damaging mutations affecting the receptors themselves or due to changes in proteins participating in the receptors’ signaling pathways. As a result, many GPCRs constitute attractive pharmacological targets for cancer management and treatment (49). A characteristic example for a type of cancer with GPCRs as potential pharmacological targets is the malignant melanoma (50). Melanoma is a type of skin cancer that develops from melanocytes, melanin-producing cells primarily located in the stratum basale, the bottom layer of the skin’s epidermis. It is one of the most dangerous types of skin cancer; while it is highly treatable if detected early. Advanced melanoma can spread to the lymph nodes and internal organs (51). A promising treatment involves the signaling pathways of endothelins, a group of peptide hormones (52–54). Endothelin signaling is conducted through two GPCRs, the endothelin receptors A and B (EDNRA and EDNRB, respectively). The two GPCRs have been implicated in various types of cancer (50) and the use of endothelin antagonists has been found to diminish cancer cells *in vitro* and *in vivo*, with both receptors having an impact (52). However, while in several types of cancer both EDNRA and EDNRB are promising drug targets, in the case of melanoma, EDNRB targeting seems to have a more profound impact upon cancer progression compared to EDNRA (54).

To analyze the signaling pathways of EDNRA and EDNRB and show the importance of the latter in melanoma, we constructed a multilayered interaction network (Figure 4). Initial interaction evidence for GPCR-related signaling in skin melanocytes was retrieved from hGPCRnet, a web application which allows the tissue-specific visualization and analysis of GPCR signal transduction networks, based on gene expression evidence on different tissues and cell types (48). The initial skin melanocyte network was then filtered to retrieve the signaling pathways of EDNRA and EDNRB, as well as all the related protein-protein interactions, which were subsequently used to construct a 3D representation in Arena3D^web^. The GPCR signaling network obtained is organized in four distinct layers: the top layer includes the receptors; namely, EDNRA and EDNRB. The two intermediate layers represent G-proteins and GRKs, for the intermediate steps of the G-protein and phosphorylation/internalization signaling pathways, respectively. Finally, the bottom layer contains all effectors that are regulated from the aforementioned pathways. Visual inspection of the EDNRA-EDNRB network (Figure 4) reveals a rather complex signaling network for the two receptors. The network consists of a total of 331 interactions; these include not only inter-layer links (e.g. GPCR-G-protein, G-protein-effector contacts), representing the different steps of the signaling pathways, but also intra-layer contacts, particularly in the effectors layer, as many effectors form supramolecular complexes with one-another to achieve a common cell response (e.g. tubulin subunits organize to microtubules). The edges representing the signaling pathways of the two receptors have been colored using red for the pathways unique to EDNRA, blue for the pathways unique to EDNRB and green for pathways controlled by both receptors.

**Figure 4.**
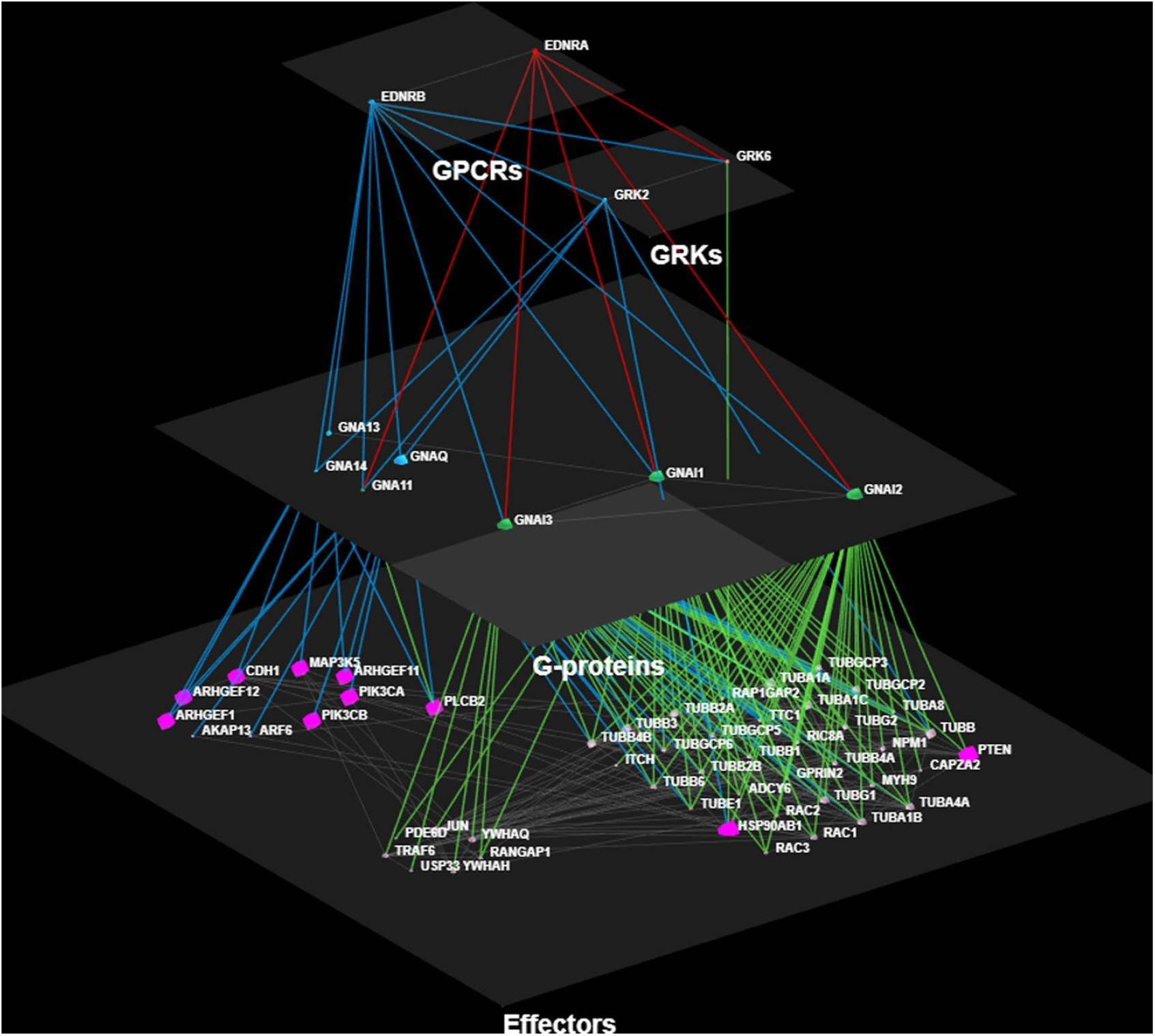
Overview of the EDNRA-EDNRB endothelin receptor signaling network. Nodes are placed in four different layers representing the different hierarchical levels in GPCR signaling (GPCRs, GRKs, G-proteins and effectors). The inter-layer edges representing the signaling pathways of the two receptors have been colored using red for the pathways unique to EDNRA, blue for the pathways unique to EDNRB and green for pathways controlled by both receptors. Intra-layer edges, connecting proteins in the same hierarchical level (e.g. effector-effector interactions) are colored grey. Effectors with clinical relevance to melanoma (CDH1, MAP3K5 etc) are shown bigger and colored magenta.

Inspection of the EDNRA-EDNRB signaling network reveals that the vast majority of endothelin-related cell responses in skin melanocytes are controlled by both receptors. Activated EDNRA and EDNRB both interact with three major G-protein subunits, GNAI1, GNAI2 and GNAI3, belonging to the Gα_i/o_ family; through these, the two GPCRs control the formation of cytoskeletal components by influencing the functionality of α-, β- and γ-tubulins (TUBA1A, TUBA1B, TUBB1, TUBG1 etc), the folding and maturation of proteins through interactions with chaperones such as HSP90-β (HSP90AB1), and the mechanisms of apoptosis and cell death through the control of apoptotic factors such as TNF-associated factor 6 (TRAF6) and the PTEN phosphatase (PTEN). Positive or negative disruption of the aforementioned processes can lead to the uncontrolled cell growth/multiplication or cell death, respectively; thus, showcasing the importance of both receptors in various types of cancer. At the same time, both receptors interact with GNA11, a member of the Gα_q_ family, which controls the function of Phospholipase C (PLCB2). The latter is an enzyme that initiates signaling cascades of second messenger molecules that regulate Protein Kinase C (PKC) and in turn, control the MAPK kinase pathway (MAP3K5). The MAPK pathway is typically deregulated in melanoma; therefore inhibitors for key members of that pathway have been used for treatment in the past (55).

However, while EDNRA’s signaling mechanisms are mostly limited to the above, a number of pathways exist that are unique to EDNRB. These feature interactions of the receptor with all members of the Gα_q/11_ family (GNAQ, GNA11, GNA13 and GNA14), which also control PLCβ and MAPK. EDNRB controls the activation of the phosphatidylinositol 3 kinases (PIKC3A and PIK3CB); these effectors play a key role in inducing drug resistance after treatment in melanoma patients and are, therefore, a decisive target for novel drug design in melanoma therapy (56). Another pathway includes the regulation of Rho Guanine Exchange Factors (RhoGEFs). EDNRB, through GNAQ and GNA13, regulates the function of RhoGEFs 1 (ARHGEF1), 11 (ARHGEF11) and 12 (ARHGEF12); these effectors regulate the enzyme activity of Rho GTPases that control the movement and morphology of melanoma cells (57). Finally, EDNRB controls the function of Cadherin-1 (CDH1), a regulator of cell adhesion and a tumor suppressor in melanoma (58). Overall, the aforementioned pathways show that while both GPCRs control a number of functions and cell mechanisms related to the progression of cancer in general and melanoma in particular, including the MAPK pathway, EDNRB exclusively regulates a number of additional signaling mechanisms that are specific to this type of cancer and is, therefore, a more suitable target for the design of anti-melanoma treatments.

## DISCUSSION

To avoid confusion, while Arena3D^web^ has been inspired by the Arena3D standalone version, it is a new application which has been written from scratch using different technologies. Compared to Arena3D, Arena3D^web^ comes with a much richer functionality and navigation options. A characteristic example is its greater variety of layout algorithms which can be used very efficiently and in multiple ways. The most advanced feature in Arena3D^web^ is the possibility to apply the layout algorithms to a subset of layers or the whole network. This allows us to generate views with much less line crossovers and makes it easier to detect feature patterns. Furthermore, as a web server application, Arena3D^web^ can run on any browser and attract a much broader audience beyond the biomedical field. On the contrary, Arena3D comes with many library dependencies, could run only on Windows and Linux and was bound to certain Java and OpenGL versions as well to the outdated Java3D library. Finally, Arena3D^web^ comes with a more flexible input file format as well as with a more advanced export function to save and re-import the status of a network at any time.

Concerning scalability, in its online version, Arena3D^web^ can handle networks of up to 5,000 nodes. This limitation can be bypassed when running the application locally, by adjusting the *max_allowed_edges* variable value in the global.R file. We tested Arena3D^web^ on a 32GB RAM, 6-core processor @ 4.2 GHz, AMD Radeon RX 5700 XT desktop machine. We ran the application οn all platforms (Unix, Windows, MacOS) using a network consisting of 1,600 nodes and 2,396 edges. We tested the application on Rstudio, Mozilla Firefox, Google Chrome, Microsoft Edge and Safari and benchmarked its memory and CPU requirements. We observed that Arena3D^web^ consumed ~300-500MB RAM on Edge browser and ~1-1.5GB RAM on Chrome, Firefox and Rstudio (including the memory needed by each browser). Due to a memory leak, on Windows systems, Mozilla Firefox kept exhausting the memory during execution, an issue which might have been evoked by a conflict between Firefox and the WebGL renderer. This problem does not occur when running Firefox on Linux. The memory requirements on MacOS across all three browsers was significantly lower (~200MB for Chrome and Safari and ~500MB for Firefox). In addition, enabling hardware acceleration on a browser helps in achieving smoother 3D transformations. Hence, a CPU load reduction from ~50% to ~13% was observed when visualizing the aforementioned network at 30 frames per seconds (FPS). Finally, the navigation controls run smoother on any of the browsers compared to RStudio, and therefore Unix and MacOS users are encouraged to run Arena3D^web^ on any of those. Windows users are encouraged to run Arena3D^web^ on Microsoft Edge which was found to be the most memory efficient.

Overall, due to its ease-of-use, simplicity and interoperability, we expect Arena3D^web^ to become the reference tool for multilayer graph analysis and visualization.

## AVAILABILITY

Arena3D^web^ is an open source application. Its code can be downloaded from the GitHub repository: https://github.com/PavlopoulosLab/Arena3Dweb. The service is available online at http://bib.fleming.gr:3838/Arena3D.

## AUTHOR CONTRIBUTIONS

EK wrote the code and implemented the tool. FB generated all the necessary data related to the case studies. NAP benchmarked the tool. RS conceived new concepts which make Arena3D^web^ more accessible than Arena3D. GAP conceived the idea and supervised the whole project from the beginning to its end. All authors wrote parts of the article.

